# Prioritizing variants in complete Hereditary Breast and Ovarian Cancer (HBOC) genes in patients lacking known *BRCA* mutations

**DOI:** 10.1101/039206

**Authors:** Natasha G. Caminsky, Eliseos J. Mucaki, Ami M. Perri, Ruipeng Lu, Joan H. M. Knoll, Peter K. Rogan

## Abstract

*BRCA1* and *BRCA2* testing for HBOC does not identify all pathogenic variants. Sequencing of 20 complete genes in HBOC patients with uninformative test results (N=287), including non-coding and flanking sequences of *ATM, BARD1, BRCA1, BRCA2, CDH1, CHEK2, EPCAM, MLH1, MRE11A, MSH2, MSH6, MUTYH, NBN, PALB2, PMS2, PTEN, RAD51B, STK11, TP53*, and *XRCC2*, identified 38,372 unique variants. We apply information theory (IT) to predict and prioritize non-coding variants of uncertain significance (VUS) in regulatory, coding, and intronic regions based on changes in binding sites in these genes. Besides mRNA splicing, IT provides a common framework to evaluate potential affinity changes in transcription factor (TFBSs), splicing regulatory (SRBSs), and RNA-binding protein (RBBSs) binding sites following mutation. We prioritized variants affecting the strengths of 10 splice sites (4 natural, 6 cryptic), 148 SRBS, 36 TFBS, and 31 RBBS. Three variants were also prioritized based on their predicted effects on mRNA secondary (2°) structure, and 17 for pseudoexon activation. Additionally, 4 frameshift, 2 in-frame deletions, and 5 stop-gain mutations were identified. When combined with pedigree information, complete gene sequence analysis can focus attention on a limited set of variants in a wide spectrum of functional mutation types for downstream functional and co-segregation analysis.

## INTRODUCTION

Currently, the lifetime risk for a woman to develop breast cancer (BC) is 12.3% and 1.3% in the case of ovarian cancer (OC [Howlander et al. 2014]). Approximately 5-10% of all BC cases are hereditary in nature, versus 25% for OC, where relative risk (RR) of BC or OC with one affected 1^st^ degree family member is estimated at 2.1 and 3.1, respectively (Stratton et al. 1998; Walsh et al. 2011). Two highly penetrant genes, *BRCA1* and *BRCA2*, are associated with a large proportion of HBOC cases. However, the estimated rate of linkage to these genes is significantly higher than the proportion of pathogenic mutations identified in HBOC families (Ford et al. 1998), suggesting unrecognized or unidentified variants in *BRCA1/2*.

Clinical *BRCA1/2* testing is restricted primarily to coding regions. Limitations on how variants can be interpreted, lack of functional validation, and mutations in other genes contribute to uninformative results. The heritability that is not associated with *BRCA* genes is likely due to other genetic factors rather than environmental causes, specifically moderate-and low-risk susceptibility genes (Antoniou and Easton 2006). Hollestelle et al. (2010) point out the challenges in estimating increased risks associated with mutations in these genes, as the disease patterns are often incompletely penetrant, and require large pedigree studies to confidently assess pathogenicity.

Next-generation sequencing (NGS) of gene panels for large cohorts of affected and unaffected individuals has become an increasingly popular approach to confront these challenges. Numerous HBOC gene variants have been catalogued, including cases in which RR has been determined; however the literature is also flooded with variants lacking a clinical interpretation (Cassa et al. 2012). It is not feasible to functionally evaluate the effects all of the VUS identified by NGS. Further, *in silico* assessment of protein coding variants has not been entirely reliable (Vihinen 2013; Rogan and Zou 2013). Several approaches have been developed to better assess variants from exome and genome-wide NGS data (Duzkale et al. 2013; Kircher et al. 2014). Nevertheless, there is an unmet need for other methods that quickly and accurately bridge variant identification and classification.

To begin to address this problem, we sought to provide potentially novel interpretations of non-coding sequence changes, based on disruption or acquisition of interactions with proteins that recognize nucleic acid binding sites. Information theory (IT)-based analysis predicts changes in sequence binding affinity, and has been applied and validated for use in the analysis of splice sites, SRBSs (Rogan et al. 1998; Rogan et al. 2003; Mucaki et al. 2013; Caminsky et al. 2015) and TFBSs (Gadiraju et al. 2003). A unified framework based on IT requires binding genome-scale site data devoid of consensus sequence bias (Schneider 1997), for example, PAR-CLIP [Photoactivatable-Ribonucleoside-Enhanced Crosslinking and Immunoprecipitation], ChIP-Seq, and a comprehensive, validated set of splice sites. Although these data sources are heterogeneous, the IT models and binding site affinities derived from them are uniformly scaled (in units of bits). Thus, binding interactions involving disparate proteins or other recognition molecules can be measured and directly compared.

We have described a unified IT framework for the identification and prioritization of variants in coding and non-coding regions of *BRCA1, BRCA2*, and 5 other HBOC genes *(ATM, CDH1, CHEK2, PALB2*, and *TP53* [Mucaki et al. submitted; biorxiv preprint: http://dx.doi.org/10.1101/031419]). This approach was applied to a cohort of 102 individuals lacking *BRCA*—mutations with a history of HBOC. This distinguished prioritized variants from flagged alleles conferring small changes to regulatory protein binding site sequences in 70.6% of cases (Mucaki et al. submitted).

In the present study, we have sequenced 13 additional genes that have been deemed HBOC susceptibility loci (*BARD1, EPCAM, MLH1, MRE11A, MSH2, MSH6, MUTYH, NBN, PMS2, PTEN, RAD51B, STK11*, and *XRCC2* [Minion et al. 2015]). These genes encode proteins with roles in DNA repair, surveillance, and cell cycle regulation (Figure 1; for further evidence supporting this gene set see **Supplementary Table S1** [Apostolou and Fostira 2013; Al Bakir and Gabra 2014]), and are associated with specific disease syndromes that confer an increased risk of BC and OC, as well as many other types of cancer (**Supplementary Table S2**). High-risk genes confer > 4-times increased risk of BC compared to the general population. *BRCA1* and *BRCA2* are estimated to increase risk 20-fold (Antoniou et al. 2003). Pathogenic variants in other high-risk genes, *CDH1, PTEN, STK11*, and *TP53*, are rarely seen outside of their associated syndromes, and account for < 1% of hereditary BC cases (Maxwell and Domchek 2013). *EPCAM, MLH1, MSH2, MSH6*, and *PMS2* have also been proposed to harbor high-risk BC alleles, but the RR is still controversial (Maxwell and Domchek 2013). Genes with moderate-risk alleles, *ATM, CHEK2*, and *PALB2*, cause between a 2- and 4-fold increased risk of BC (Apostolou and Fostira 2013; Maxwell and Domchek 2013). The remaining genes (*BARD1, MRE11A, MUTYH, NBN, RAD51B*, and *XRCC2)* are newly identified and currently associated with unknown risks for HBOC (Figure 1).

**Figure 1.**
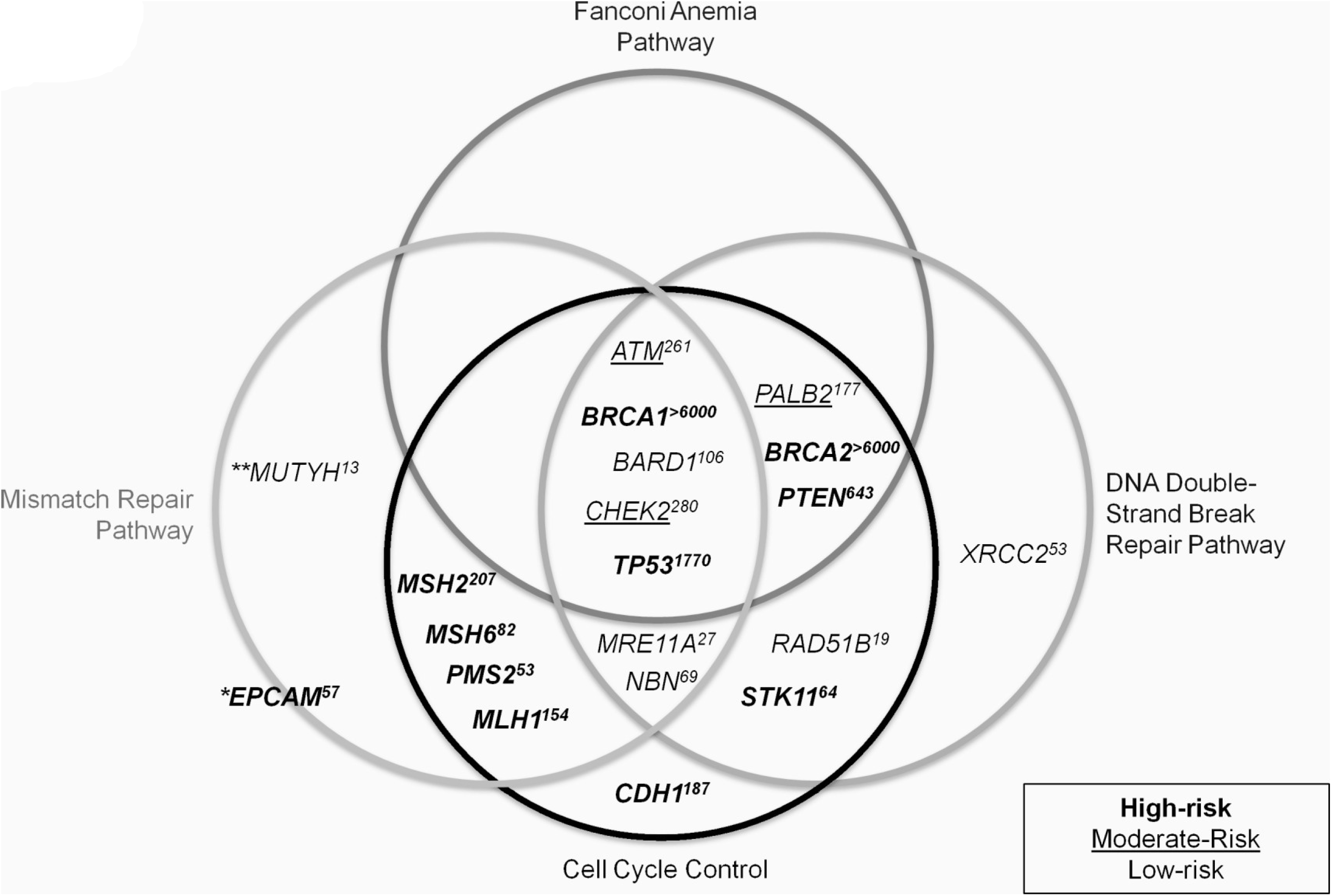
Common Genomic Pathways Among 20 HBOC Genes, Including Risk and Relevant Literature. The left, top, and right circles indicate sequenced genes that play important roles in the MMR, Fanconi Anemia, and DNA double-strand break repair pathways, respectively. The bottom circle contains genes involved in cell cycle control. Genes considered to present a high risk of breast and/or ovarian cancer when mutated are bolded, moderate-risk genes are underlined, and low-risk genes are in normal font. The estimated number of articles listing a gene’s association with breast or ovarian cancer (based on a systematic search in PubMed [performed June 2015]) is indicated in superscript. *\*MUTYH* is only high risk in the case of bi-allelic mutations. *\*\*EPCAM* is not involved in any pathways, but is associated with hereditary non-polyposis colorectal cancer (HNPCC) by virtue of the fact that 3 ‘ deletions of *EPCAM* can cause epigenetic silencing of *MSH2*, causing Lynch syndrome protein. See **Supplementary Table S1** for citations and further evidence supporting this gene set.

We report NGS of hybridization-enriched, complete genic and surrounding regions of these genes, followed by variant analysis in 287 consented patients from Southwestern Ontario, Canada with previously uninformative HBOC test results (this set of patients is different from our submitted study, except for 6 previously anonymous individuals who subsequently consented to participate). We then reduced the set of potentially pathogenic gene variants in each individual by prioritizing the results of coding and IT-analyses. After applying a frequency-based filter, the IT-based framework prioritizes variants based on their predicted effect on the recognition of sequence elements involved in mRNA splicing, transcription, and untranslated region (UTR) binding, combined with UTR secondary structure and coding variant analysis. Our approach integrates disparate sources of information, including bioinformatic analyses, likelihood ratios based on familial segregation, allele frequencies, and published findings to prioritize disease-associated mutation candidates.

## METHODS

### Ethics and Patient Recruitment

Recruitment and consent of human participants was approved by the University of Western Ontario Research Ethics Board (Protocol 103746). Patients were enrolled from January, 2014 through March, 2015 at London Health Sciences Centre (LHSC). Patients met the following criteria: male or female, aged between 25 and 75 years, > 10% risk of having an inherited mutation in a breast/ovarian cancer gene, diagnosed with BC and/or OC, and previously receiving uninformative results for a known, pathogenic *BRCA1* or *BRCA2* variant in either the patient or other relatives (by Protein Truncation Test [PTT], Denaturing High performance Chromatography [DHPLC], and/or Multiplex Ligation-dependent Probe Amplification [MLPA]).

The median age of onset for patients (N=287; **Supplementary Figure S1**) with BC was 48 (N=277), and 46 for OC (N=17), and 7 were diagnosed with both BC and OC. Furthermore, 31 patients had bilateral BC (98 patients at diagnosis; 23 developed tumors on the opposite side after the initial occurrence), 1 had bilateral OC, and 13 have had recurrent BC in the same breast. There was a single case of male BC (**Supplementary Table S3**).

### Probe Design, Sample Preparation, and Sequencing

Probes for sequence capture were designed by *ab initio* single copy analysis, as described in Mucaki et al. (submitted) and Dorman et al. (2013). The probes covered 1,103,029 nt across the 21 sequenced genes, including the negative control gene *ATP8B1* (see **Supplementary Methods** for gene names, GenBank accession numbers, and OMIM reference numbers). This set of genes was proposed for evaluation at the Evidence-based Network for the Interpretation of Germline Mutant Alleles (ENIGMA) Consortium Meeting (2013). Other genes that have been found to be mutated in HBOC could not be included (eg. *BRIP1, RAD50, RAD51C, RAD51D* [Heikkinen et al. 2003; Seal et al. 2006; Janatova et al. 2015]).

Patient DNA extracted from peripheral blood was either obtained from the initial genetic testing at LHSC Molecular Genetics Laboratory or isolated from recent samples. NGS libraries were prepared using modifications to a published protocol (Gnirke et al. 2009) described in Mucaki et al. (submitted), and all post-capture pull-down steps were automated (**Supplementary Methods**). An Illumina Genome Analyzer IIx instrument in our laboratory was used for sequencing.

Library preparation and re-sequencing were repeated for samples with initial average coverage below our minimum threshold (< 30x). To ensure that the proper sample was re-sequenced, the variant call format (VCF) files from each run were compared to all others in the run using VCF-compare (http://vcftools.sourceforge.net/). VCF files from separate runs for the re-sequenced patient were concordant, except for minor differences in variant call rates due to differences in coverage. The aligned reads from both runs were then merged (with BAMtools; http://sourceforge.net/projects/bamtools/).

Samples were demultiplexed and aligned using CASAVA (Consensus Assessment of Sequencing and Variation; v1.8.2; DePristo et al. 2011) and CRAC (Complex Reads Analysis & Classification; v1.3.0; http://crac.gforge.inria.fr/). Aligned BAM files were then pre-processed for variant calling with Picard (v.1.109; http://broadinstitute.github.io/picard/) (MarkDuplicates, AddorReplaceReadGroups, FixMateInformation). The Genome Analysis Toolkit (GATK; v3.1; http://www.broadinstitute.org/gatk/) was then used for variant calling using the modules ‘Indel realigner’ and the ‘Unified Genotyper’. Variants flagged by bioinformatic analysis [see *Variant Analysis* below] were also assessed by manual inspection with the Integrative Genome Viewer v2.3 (IGV; http://www.broadinstitute.org/igv/). Variants in this study are written in HGVS notation, are based on cDNA sequence, and comply with journal guidelines.

### Information Models

Models for natural splice sites (SSs) and splicing regulatory factors (SRFs) are described in Mucaki et al. (2013). These models were used to predict deleterious effects on natural splicing, the activation of cryptic SSs, and changes to binding of splicing enhancers and silencers. In addition, using a combination of cryptic site activation and hnRNPA1 site prediction, pseudoexon formation was also assessed.

We previously built models for TFBSs (N=83) using ENCODE ChIP-seq data (ENCODE Project Consortium 2012; Mucaki et al. submitted). Due to the inclusion of the additional genes, 8 additional transcription factors (TFs) were identified from published literature and ENCODE data from breast cancer cell lines to exhibit ChIP-seq evidence of binding and potentially, regulating these genes. However, models for 3 of these TFs passed our quality control criteria (TFIIIB150 [*BDP1*], PBX3 and ZNF274; described in Lu et al. submitted). **Supplementary Table S4** contains the full list of TFs (N=86) and indicates which genes exhibit evidence of promoter or other binding events. Noise models (N=5), reflecting motifs of interacting cofactors or sequence-specific histone modifying events, were excluded (**Supplementary Methods**).

Information weight matrices, R_i_(*b,l*), for sequences bound by RNA-binding proteins (RBPs) were derived from frequency matrices published in the Catalog of Inferred Sequence Binding Preferences of RNA binding protein (CISBP-RNA; http://cisbp-rna.ccbr.utoronto.ca/) and RNA Binding Protein Database (BPDB; http://rbpdb.ccbr.utoronto.ca/). These R_i_(*b,l*)’s were used to compute changes in binding affinity due to SNVs, using conservative minimum information thresholds described in Mucaki et al. (submitted). Finally, predicted changes in UTR structure resulting from variants were determined using SNPfold (http://ribosnitch.bio.unc.edu/snpfold/; Halvorsen et al. 2010). Significant changes in UTR structure and stability were represented using mfold (http://unafold.rna.albany.edu/?q=mfold).

### Variant Analysis

Information analysis has been used in the interpretation of variant effects on binding sites containing these changes, whether this involves the creation or strengthening, or the abolition or weakening of a site (Rogan et al. 1998). This analysis was applied to all variants identified by NGS. Changes in information are directly related to changes in thermodynamic entropy and thus binding affinity (Rogan et al. 1998). For example, a 1.0 bit change in information corresponds to at least a 2-fold change in binding affinity. Information theoretical analysis of SSs and SRF binding sites has been extensively used and proven to be reliable and robust (85.2% accuracy when compared to variants validated by expression studies) (Caminsky et al. 2015).

Information analysis was automated and thresholds for changes were applied programmatically based on our previously validated criteria (Rogan et al. 1998; Rogan et al. 2003; von Kodolitsch et al. 2006; Dorman et al. 2014). This reduced manual review of prioritized variants, databases and the literature. A minimum 1.0 bit threshold was set for variants predicted to affect natural splice sites or that activate a cryptic splice site by exceeding the strength of cognate natural sites. Variants affecting splicing regulatory, transcription, and RNA-binding protein binding sites were assessed more stringently and had a minimum threshold of 4.0 bits, i.e. ≥16 fold, in order to be flagged for further assessment. A population frequency filter was also applied to variants with allele frequencies > 1% (in dbSNP) or > 5% of our patient cohort, which were eliminated from further consideration.

To assess coding changes affecting predicted protein chain length or amino acid(s) composition, we used SNPnexus (http://snp-nexus.org/). Insertion/deletions (indels) and nonsense mutations were noted, and missense variants were further assessed with *in silico* tools (Mutation Assessor – http://mutationassessor.org/, PolyPhen2 - http://genetics.bwh.harvard.edu/pph2/, PROVEAN/SIFT – http://provean.jcvi.org/), by referencing the published literature, and consulting mutation databases [listed in **Supplementary Table S5;** see Mucaki et al. (submitted) for more details on variant analysis]. Variants remained prioritized unless there was clear evidence (co-segregation analysis or functional assays) supporting the non-pathogenicity of the variant.

*EPCAM* mutations in familial cancer are limited to 3’ deletions causing epigenetic silencing of *MSH2*, and there is currently no evidence of other types of variants that alter its mRNA transcript or protein product (Ligtenberg et al. 2009). Therefore, with the exception of indels, none of the variants flagged in *EPCAM* were prioritized. We chose to prioritize variants in *MUTYH* using the same framework as all other genes, despite *MUTYH* pathogenicity resulting from biallelic variants (Jones et al. 2002), because it is possible that a second *MUTYH* mutation remains unrecognized.

All protein truncating (nonsense and indels) as well as potentially pathogenic splicing and missense mutations were Sanger sequenced for confirmation (details in **Supplementary Table S6**).

#### Negative Control

Variants present in the *ATP8B1* gene were used as negative controls for our variant analysis framework. Initially, it was included in the list of prioritized HBOC genes provided by ENIGMA, but evidence for its association with HBOC is lacking in the published literature. Furthermore, it is not a known susceptibility gene for any type of cancer (mutations in *ATP8B1* cause progressive familial intrahepatic cholestasis [Gonzales et al. 2014]), and is infrequently mutated in breast tumors in several studies (for example, see Cancer Genome Atlas Network [2012]).

### Likelihood Ratios (LRs)

Patients with prioritized coding and/or splicing variants, which we consider the most likely to be pathogenic, were selected for co-segregation analysis (N=24) using an online tool that calculates the likelihood of a variant being deleterious based on pedigree information (https://www.msbi.nl/cosegregation/; Mohammadi et al. [2009]). Genotypes were assigned based on phenotype such that family members with breast or ovarian cancer at any age were assigned the same genotype as the patient in our study (“carrier”) and family members affected by other cancers, other diseases, or who are disease-free were assigned the “non-carrier” genotype. Because the penetrance parameters cannot be altered from the settings given for *BRCA1* or *BRCA2*, the *BRCA2* option was selected for patients with prioritized variants in non*-BRCA* genes. Penetrance in *BRCA2* is known to be lower than *BRCA1* values (Mohammadi et al. 2009). Current evidence suggests that mutations in non*-BRCA* genes may be less penetrant than those in the *BRCA* genes (Apostolou and Fostira 2013), however the penetrance of many of these variants remains unknown (**Supplementary Methods**).

## RESULTS

### Variant Analysis

We identified 38,372 unique variants among 287 patients (26,636 intronic, 7,287 intergenic, and 714 coding), on average 1,975 variants per patient, before any filtering criteria were applied. The extensive span of sequences captured in this study, i.e. complete genes and flanking regions, constrained the genomic density and sequence coverage that could be achieved; this precluded accurate copy number estimation based solely on read counts.

#### Natural Site Variants

The Shannon Human Splicing Mutation Pipeline (http://www.mutationforecaster.com; Shirley et al. [2013]) was used to predict the effect of the 14,458 variants that could potentially affect splicing, of which 244 reduced natural SS strength. Further stringent filtering of the natural SS based on information content changes and allele frequency resulted in 7 flagged variants (**Supplementary Table S7**). Henceforth, allele frequency of known variants can be found in their associated supplemental table (where available).

Four of these variants were prioritized (Table 1). A novel synonymous variant in exon 2 of *RAD51B*, c.84G>A (p.Gln28=), is predicted to increase exon skipping by weakening the natural splice donor (*R_i, final_* = 5.2 bits, Δ*R_i_* = −3.0 bits). A known *ATM* variant, c.6198+1G>A (8-1D.9-1B [Stankovic et al. 1998]), abolishes the natural donor SS of constitutively spliced exon 42 (*R_i, final_* = −13.7 bits, Δ*R_i_* = −18.6 bits). There is no evidence in public databases for appreciable alternative splicing of this exon in normal breast tissues. The variant will either lead to exon skipping or activation of a pre-existing cryptic site (Figure 2). An Ataxia-Telangiectasia patient with this variant exhibited low expression, protein truncation, and abolished kinase activity of ATM (Reiman et al. 2011). *MLH1* c.306+4A>G causes increased exon skipping (and a decrease in wild-type exon relative expression) due to the weakening (*R_i, final_* = 6.0 bits, Δ*R_i_* = −2.6 bits) of the exon 3 natural donor. Tournier et al. (2008) assessed this variant using an *ex vivo* splicing assay and observed cryptic site activation and exon 3 skipping. *MRE11A* c.2070+2A>T is indicated in ClinVar as likely pathogenic and abolishes the natural donor site of exon 19 (*R_i, final_* = −11.0 bits, Δ*R_i_* = −18.6 bits), while strengthening a cryptic site 5 nt upstream of the splice junction (*R_i, final_* = 8.1 bits, Δ*R_i_* = 0.6 bits). Either cryptic SS activation or complete exon skipping are predicted.

**Table 1.** Prioritized Variants Predicted by IT to Affect Natural and Cryptic Splicing

| Gene | Variant | rsID (dbSNP142)<br>Allele Frequency (%)<sup>†</sup> | Information Change |  |  | Consequence |
| --- | --- | --- | --- | --- | --- | --- |
| | | | $R_{i,initial}$<br>(bits) | $R_{i,final}$<br>(bits) | $\Delta R_i$<br>(bits) | |
| *ATM* | NM_000051.3:c.6198+1G>A<sup>1,2</sup> | - | 4.9 | -13.7 | -18.6 | Abolished natural<sup>a,d</sup> |
| *MRE11A* | NM_005591.3:c.2070+2A>T* | - | 7.6 | -11 | -18.6 | Abolished natural<sup>a,d</sup> |
| *MLH1* | NM_000249.2:c.306+4A>G*<sup>3</sup> | rs267607733 | 8.6 | 6 | -2.6 | Weakened natural<sup>b</sup> |
| *RAD51B* | NM_002877.4:c.84G>A*<br>p.Gln28= | Novel | 8.2 | 5.2 | -3 | Weakened natural<sup>a</sup> |
| *BARD1* | NM_000465.2:c.1454C>T*<br>p.Ala485Val | Novel | -2.7 | 4.4 | 7.1 | Created cryptic<sup>b</sup> |
| *BRCA1* | NM_007294.2:c.5074+107C>T | rs373676607 | -1.3 | 5.7 | 7 | Created cryptic<sup>c,e</sup> |
| *CDH1* | NM_004360.3:c.1223C>G*<sup>4</sup><br>p.Ala408Gly | Novel | -0.6 | 4.3 | 4.9 | Created cryptic<sup>b</sup> |
| *RAD51B* | NM_002877.4:c.958-29A>T*** | rs34436700<br>0.78 | 2.2 | 4.4 | 2.2 | Strengthened cryptic<sup>c</sup> |
| *STK11* | NM_000455.4:c.375-194GT>AC | rs35113943 17.61<br>rs117211142 0.80 | 7.5 | 8.8 | 1.3 | Strengthened cryptic<sup>c</sup> |
| *XRCC2* | NM_005431.1:c.122-154G>T | Novel | 8.1 | 10 | 1.9 | Strengthened cryptic<sup>c</sup> |

\*Confirmed by Sanger sequencing; \*\*\*Ambiguous Sanger sequencing results; <sup>†</sup>If available; <sup>a</sup>exon skipping; <sup>b</sup>exon truncation; <sup>c</sup>intron retention; <sup>d</sup>use of alternate isoform; <sup>e</sup>reduced expression of natural isoform; <sup>1</sup>Stankovic et al., 1998, <sup>2</sup>Reiman et al., 2011, <sup>3</sup>Tournier et al., 2008, <sup>4</sup>Schrader et al., 2011.

**Figure 2.**
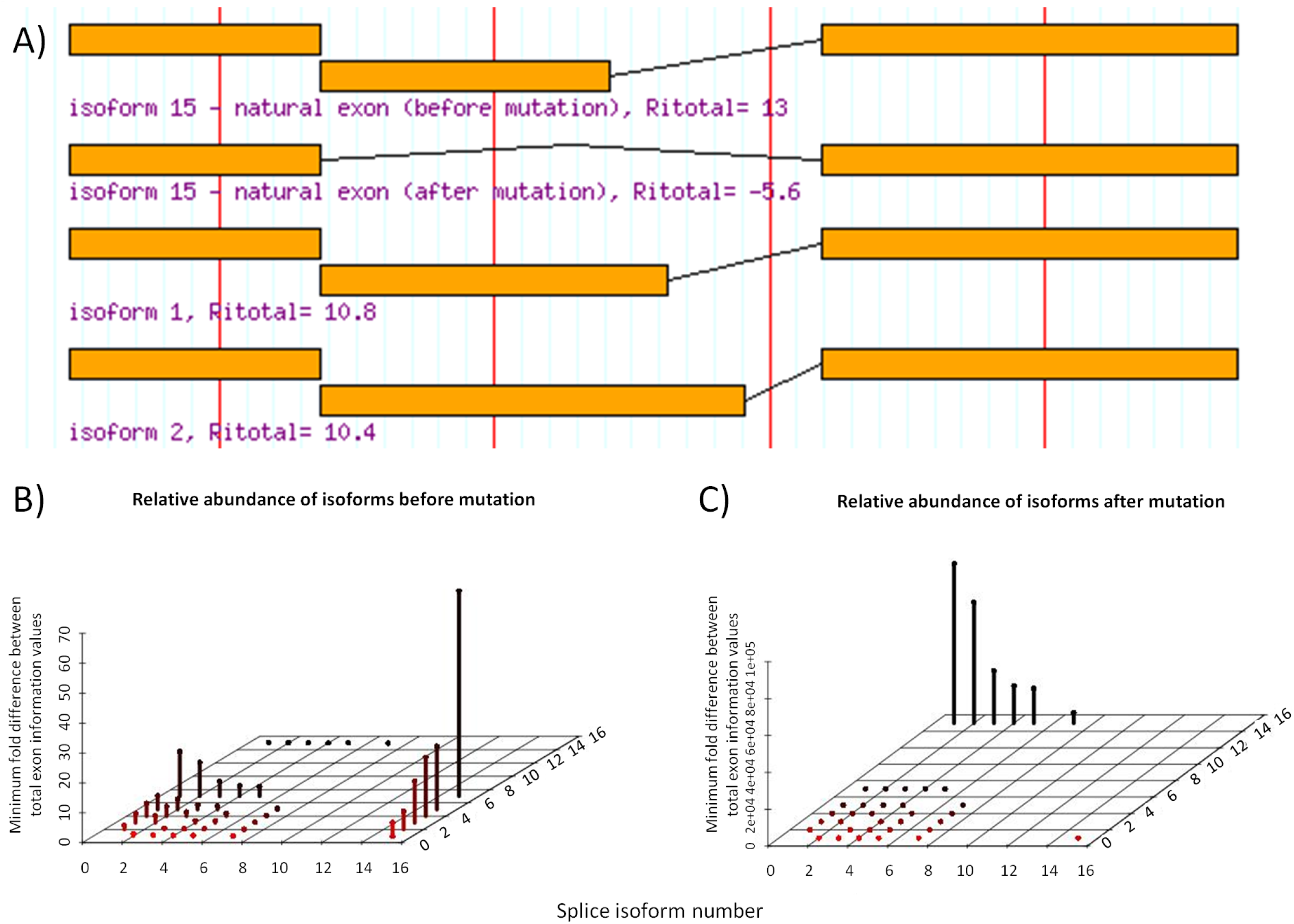
Predicted Isoforms and Relative Abundance as a Consequence of *ATM* natural splice variant c.6198+1G>A. **A**) Intronic ATM variant c.6198+1G>A abolishes the natural donor of exon 42 (*R_i, initial_* = 4.9 bits, *ΔR_i_* = −18.6 bits), and would either result in exon skipping (causing a frame-shift; isoform 15 after mutation), or possibly activate a downstream cryptic site (isoform 1 maintains reading frame, isoform 2 would not). **B**) The peaks in plot display the predicted abundance (Y-axis) of a splice isoform (X-axis) relative to another predicted isoform (Z-axis). In the wild type mRNA, the natural exon (isoform 15) has the highest predicted relative abundance. Before mutation, it is predicted to be ∽5 fold stronger than isoform 1 and 2. **C**) After mutation, isoform 1 and 2 is now > 100000 fold stronger than isoform 15 (abolished wildtype exon). Isoform 2 to be slightly less abundant than 1.

The *BRCA2* variant c.68-7T>A was not prioritized, as its pathogenicity has not been proven. While there is evidence that this variant induces (in-frame) exon skipping (Théry et al. 2011), it did not segregate with disease in HBOC pedigrees, where abnormal splicing was not seen (Santos et al. 2014). The *ATM* variant c.1066-6T>G, previously reported in Mucaki et al. (submitted), was also not prioritized as the variant does not correlate with breast cancer risk (Ding et al, 2011).

#### Activation of Cryptic Splicing

The Shannon Pipeline identified 9,480 variants that increased the strength of at least one cryptic site, of which 9 met or exceeded the defined thresholds for information change. Six of these were prioritized (Table 1). A novel *BARD1* variant in exon 6 (c.1454C>T; p.Ala485Val) creates a donor SS (*R_i, final_* = 4.4 bits, Δ*R_i_* = 7.1 bits), which would produce a 58 nt frameshifted exon if activated. The natural donor SS of exon 6, 116 nt downstream of the variant, is stronger (5.5 bits), but the Automated Splice Site and Exon Definition Analysis (ASSEDA http://mutationforecaster.com) server predicts equal levels of expression of both natural and cryptic exons. A *BRCA1* mutation 5074+107C>T downstream of exon 16 is predicted to extend the exon by 105 nt, and be slightly more abundant than the natural exon (*R_i, total_* of 8.6 and 8.1 bits, respectively). *CDH1* c.1223C>G (p.Ala408Gly), previously reported in a *BRCA*-negative lobular BC patient with no family history of gastric cancer (Schrader et al. 2011), creates a cryptic donor site (*R_i, final_* = 4.3 bits, Δ*R_i_* = 4.9 bits) in exon 9, 97 nt downstream of the natural acceptor. While residual splicing of the normal exon is still expected, the cryptic is predicted to become the predominant splice form (∽twice as abundant).

*STK11* c.375-194GT>AC (rs35113943 & rs117211142), and the novel *XRCC2* c.122-154G>T both strengthen strong pre-existing cryptic sites exceeding the *R_i, total_* values of their respective natural exons. Finally, a known *RAD51B* variant 29 nt upstream of exon 10: c.958-29A>T strengthens a cryptic acceptor (*R_i, final_* = 4.4 bits, Δ*R_i_* = 2.2 bits) that, if activated, would produce a transcript retaining 21 intronic nucleotides.

The remaining cryptic site variants (**Supplementary Table S7**) were not prioritized. The novel *BRCA2* c.7618-269_7618-260del10 variant is predicted to create a cryptic site with an exon having a lower *R_i, total_* value (5.2 bits) than the natural exon (6.6 bits). *PMS2* c.1688G>T (p.Arg563Leu; rs63750668; 3 patients) does not segregate with disease. Drost et al. (2013) demonstrated that this variant does not impair DNA repair activity. Finally, *RAD51B* c.728A>G (p.Lys243Arg; rs34594234; 7 patients) predicts an increase in the abundance of the cryptic exon; however the natural exon remains the predominant isoform.

#### Pseudoexon Activation

Pseudoexons arise from creation or strengthening of an intronic cryptic SS in close proximity to another intron site of opposite polarity. Our analysis detected 623 variants with such intronic cryptic sites, of which 17 were prioritized (among 9 genes), occurring within 250 nt of a preexisting site of opposite polarity, with an hnRNPA1 site within 5 nt of the acceptor of the predicted pseudoexon (**Supplementary Table S8**). Three are novel (*BRCA2* c.7007+824C>T, *BRCA2* c.8332-1130G>T, and *PTEN* c.802-796C>A), and the remainder were present in dbSNP. Seven of these variants (*BARD1* c.1315-168C>T, *BRCA2* c.631+271A>G, *MLH1* c.1559-1732A>T, *MRE11A* c.1783+2259A>G, *MSH6* c.260+1758G>A, *PTEN* c.79+4780C>T, and *RAD51B* c.1037-1012C>A), although rare, occur in multiple patients, and one patient had predicted pseudoexons in both *BARD1* and *RAD51B*.

#### SRF Binding

Variants within exons or within 500 nt of a natural SS (N=9,998) were assessed for their potential effect on SRF binding sites (SRFBSs). Initially 216 unique variants were flagged (**Supplementary Table S9),** but after considering each in the context of the SRF function and location within the gene (Caminsky et al. 2015), we prioritized 148, of which 57 are novel. Some prioritized variants affect distant SRFs that may activate cryptic sites, but were not predicted to affect natural splicing. Of the 88 suitable prioritized variants for which exon definition analysis was performed (where initial *R_i, total_* of the exon > SRF gap surprisal value), 55 were predicted to induce or contribute to increased exon skipping. For example, an uncommon *ATM* missense variant within exon 41, c.6067G>A (p.Gly2023Arg; rs11212587), strengthens an hnRNPA1 site (*R_i, final_* = 5.2 bits, Δ*R_i_* = 4.7 bits) 30 nt from the natural donor, and is predicted to induce exon 41 skipping (Δ*R_i, total_* = −9.5 bits).

#### TF binding

To assess potential changes to TFBSs, variants occurring from 10 kb upstream of the start of transcription through the end of the first intron were analyzed by IT, flagging 88 (of 4,530 identified; **Supplementary Table S10**). Considering the gene context of each TFBS and extent of information change, we prioritized 36 variants. The following illustrates the rationale for highlighting these variants: *BRCA1* c.-19-433A>G abolishes a binding site for HSF 1 (*R_i, initial_* = 5.5 bits, Δ*R_i_* = −7.8 bits). While HSF 1 is known to be a transcriptional activator associated with poor BC prognosis (Santagata et al. 2011), the specific effect of reduced HSF 1 binding to *BRCA1* has not been established. Similarly, *MLH1* c.-4285T>C (rs115211110; 5 patients) significantly weakens a C/EBPβ site (*R_i, initial_* = 10.1 bits, Δ*R_i_* = −6.3 bits), a TF that has been shown to play a role in BC development and progression (Zahnow 2009). Another *MLH1* variant, c.-6585T>C (novel), greatly decreases the binding strength (*R_i, initial_* = 12.5 bits, Δ*R_i_* = −10.8 bits) of the NF-κB p65 subunit, which is activated in ER-negative breast tumors (Biswas et al. 2004). Two prioritized variants (*PMS2* c.-9059G>C and *XRCC2* c.-163C>A) weaken PAX5 binding sites, a TF which when overexpressed, can result in mammary carcinoma cells regaining epithelial cell characteristics (Vidal et al. 2010).

#### Alterations to mRNA Structure

A total of 1,355 variants were identified in the 5’ and 3’ UTRs of the patients. Analysis of these variants with SNPfold flagged 3 unique variants (p < 0.05), in *BRCA1, BARD1*, and *XRCC2* (Table 2). The predicted mRNA 2° structures of the reference and variant sequences are shown in Figure 3 (generated with mfold). The *BRCA1* variant occurs in the 3’UTR of all known transcript isoforms (NM_007294.3:c.*1332T>C; rs8176320; 3 patients). The most likely inferred structure consisting of a short arm and a larger stem loop is destabilized when the variant nucleotide is present (Figure 3A and B). The *BARD1* variant falls within the 5’ UTR of a rare isoform (XM_005246728.1:c.-53G>T; rs143914387; 5 patients), and is within the coding region of a more common transcript (NM_000465.2:c.33G>T; p.Gln11His). While the top ranked isoform following mutation is similar to the wild-type structure, the second-ranked isoform (ΔG = +1.88kcal/mol) is distinctly different, creating a loop in a long double-stranded structure (Figure 3C and D). The *XRCC2* variant is within its common 5’ UTR (NM_005431.1:c.-76C>T) and is located 11 nt downstream from the 5’ end of the mRNA. The variant nucleotide disrupts a potential GC base pair, leading to a large stem-loop that could allow access for binding of several RBPs (Figure 3E and F). The variant simultaneously strengthens a PUM2 (*R_i, initial_* = 2.8 bits, Δ*R_i_* = 4.4 bits) and a RBM28 site (*R_i, initial_* = 4.0, Δ*R_i_* = 3.6 bits), however there is a stronger NCL site (8.3 bits) in the area that is not affected and may compete for binding.

**Table 2.** Variants Predicted by SNPfold to Significantly Affect UTR Structure

| Gene | Variant | UTR Position | rsID (dbSNP142)<br>Allele Frequency (%)<sup>†</sup> | Rank | *p*-value |
| --- | --- | --- | --- | --- | --- |
| *BARD1* | XM_005246728.1:c.-53G>T<br>(c.33G>T p.Gln11His) | 5'UTR | rs143914387<br>0.04 | 6/600 | 0.01 |
| *BRCA1* | NM_007294.3:c.*1332T>C<br>NM_007299.3:c.*1438T>C | 3'UTR | rs8176320<br>0.42 | 13/450 | 0.03 |
| *XRCC2* | NM_005431.1:c.-76C>T | 5'UTR | rs547538731<br>0.08 | 3/300 | 0.01 |

<sup>†</sup>If known

**Figure 3.**
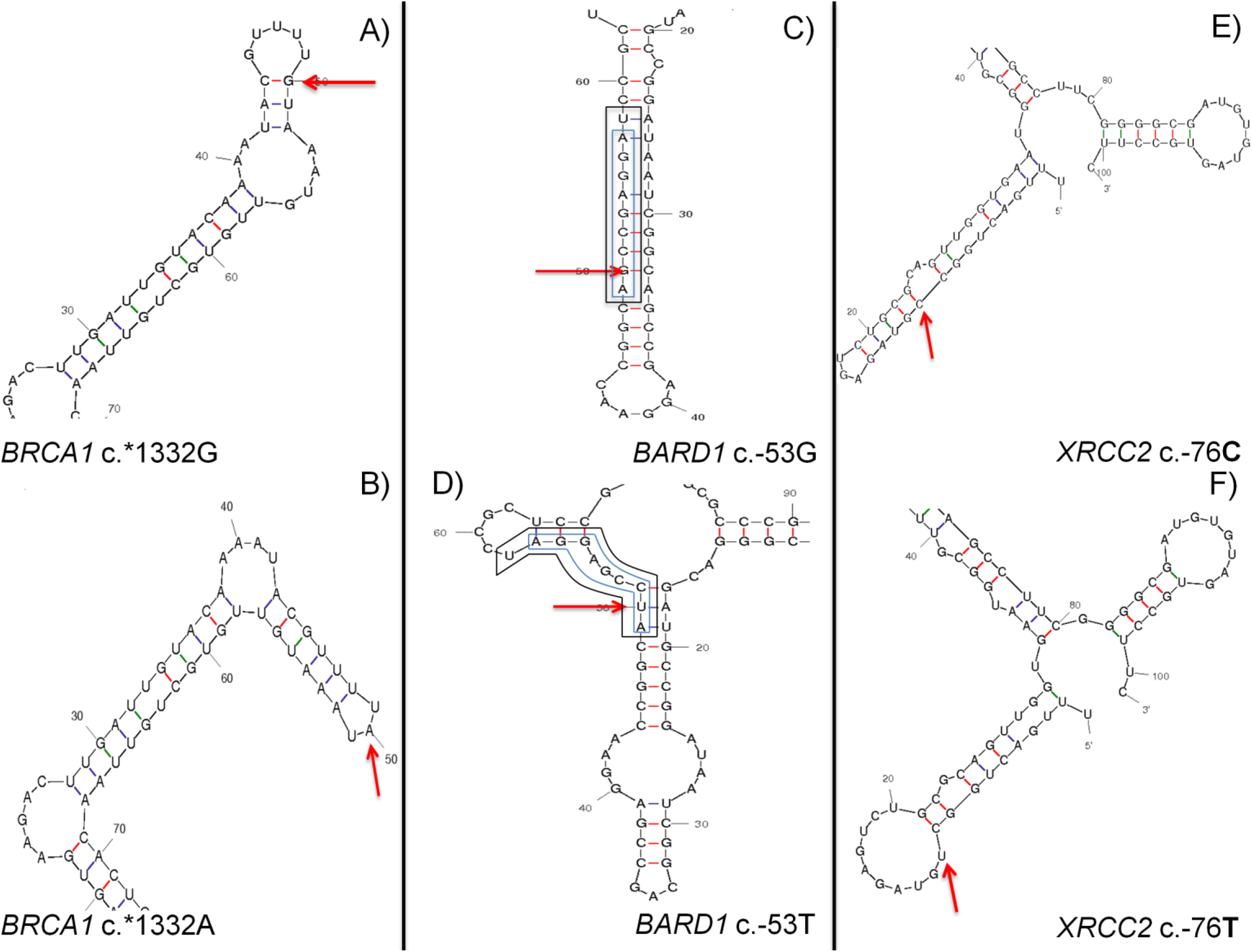
Predicted RNA Structure Change due to Variants Flagged by SNPfold using mfold. Wild-type (A, C, and E) and variant (B, D and F) structures are displayed. The variant nucleotide is marked with an arrow. **A**) Predicted wild-type structure of *BRCA1* 3’UTR surrounding c.*1332G>A. **B**) *BRCA1* 3’UTR structure due to c.*1332A variant, extending arm length while reducing hairpin size. **C**) *BARD1* 5’UTR structure of rare isoform (XM_005246728.1:c.-53G>T). Two overlapping pre-existing RBP sites (SRSF7 [outer box] and SRSF2 [inner box]) are predicted and either could occupy this location if accessible. **D**) 2° *BARD1* 5’ UTR structure of the region predicted only with sequence containing the c.-53T mutation. The primary predicted c.-53T structure is identical to wild-type (with one disrupted C-G bond leading to a 4.1 kcal/mol lower ΔG). The variant both weakens and abolishes the pre-existing SRSF7 and SRSF2 sites, respectively. **E**)*XRCC2* structure within common 5’UTR surrounding c.-76C>T variant. **F**) *XRCC2* 5’UTR structure predicted from c.-76T sequence, containing a hairpin not found in wild-type. This hairpin may allow for the binding of previously inaccessible nucleotides including the altered nucleotide.

#### RBP binding

Using IT models of 76 RBBSs, 33 UTR variants were prioritized (**Supplementary Table S11**) from the initial list of 1,367 UTR variants. Interestingly, one of the three variants that destabilized the mRNA was also flagged using our RBP scan. The *BARD1* c.-53A>C variant weakens a predicted 8.3 bit SRSF7 site (Δ*R_i_* = −3.0 bits) while simultaneously abolishing a predicted 9.7 bit SRSF2 site (Δ*R_i_* = −29.7 bits) (Figure 3C-D).

### Exonic Protein-Altering Variants

#### Protein Truncating

Of the 714 identified coding variants, 6 were indels, each of which found in a single patient, and 2 preserve the reading frame. Two indels were novel (BRCA1:c.3550_3551insA [p.Gly1184Glufs] and CDH1:c.30_32delGCT [p.Leu11del]). Previously reported indels were detected in *CHEK2* and *PALB2*. In addition, 5 nonsense mutations, which have been previously reported by others, were found in 6 different patients (Table 3; details in **Supplementary Table S12**).

**Table 3.** Variants Resulting in Premature Protein Truncation

| Gene | Exon | Variant | rsID (dbSNP142)<br>Allele Frequency (%)<sup>†</sup> | Details |
| --- | --- | --- | --- | --- |
| Frameshift Insertions/Deletions |  |  |  |  |
| *BRCA1* | 10 of 23 | NM_007294.2:c.3550_3551insA**<br>p.Gly1184Glufs | Novel | STOP at p.1187<br>676 AA short |
| *PALB2* | 4 of 13 | NM_024675.3:c.757_758delCT*<br>p.Leu253Ilefs | rs180177092 | STOP at p.255<br>932 AA short |
| *PALB2* | 9 of 13 | NM_024675.3:c.2920_2921delAA*<br>p.Lys974Glufs | rs180177126 | STOP at p.979<br>208AA short |
| Insertions/Deletions with Conserved Reading Frame |  |  |  |  |
| *CDH1* | 1 of 16 | NM_004360.3:c.30_32delGCT***<br>p.Leu11del | Novel | Loss of 1 AA<br>Frame and AA sequence conserved |
| *CHEK2* | 4 of 14 | NM_007194.3:c.483_485delAGA*<br>p.Glu161del | - | Loss of 1 AA<br>Frame and AA sequence conserved |
| Stop Codons |  |  |  |  |
| *ATM* | 13 of 63 | NM_000051.3:c.1924G>T*<br>p.Glu642Ter | - | 2415 AA short |
| *ATM* | 62 of 63 | NM_000051.3:c.8977C>T*<br>p.Arg2993Ter | - | 64 AA short |
| *BRCA1* | 23 of 23 | NM_007294.2:c.5503C>T**<br>p.Arg1835Ter | rs41293465 | 28 AA short |
| *PALB2* | 13 of 13 | NM_024675.3: c.3549C>G*<br>p.Tyr1183Ter | rs118203998 | 4 AA short |

\*Confirmed by Sanger sequencing; \*\*Not confirmed through Sanger sequencing; \*\*\*Ambiguous Sanger sequencing results; <sup>†</sup>If known; AA: amino acid

#### Missense Variants

Of the 155 unique missense variants (**Supplementary Table S13),** 119 were prioritized by consulting published literature, disease-and gene-specific databases. All are of unknown clinical significance and 21 have not been previously reported.

Missense variants that have been previously described as detrimental include the *ATM* variant c.7271T>G (p.Val2424Gly; rs28904921; 2 patients), which replaces a hydrophobic residue by glycine in the conserved FAT domain and confers a 9-fold increase (95% CI) in BC risk (Goldgar et al. 2011). Functional studies, assessing ATM kinase activity *in vitro* with TP53 as a substrate, showed that cell lines heterozygous for the mutation had less than 10% of wild-type kinase activity, such that this variant is expected to act in a dominant-negative manner (Chenevix-Trench et al. 2002). The *CHEK2* variant c.433C>T (p.Arg145Trp; rs137853007; 1 patient) results in rapid degradation of the mutant protein (Lee et al. 2001). Finally, the *PMS2* variant c.2T>C (p.Met1Thr) is listed in ClinVar as pathogenic and would be expected to abrogate correct initiation of translation. This variant has not been reported in BC families, but is associated with colorectal cancer (CRC) (Senter et al. 2008).

### Variant Prioritization

We prioritized an average of 18.2 variants in each gene, ranging from 7 (*XRCC2)* to 61 (*ATM)*, an average of 0.41 variants/kb, and an average of 0.65 variants/patient (Table 4). *ATM* had the second greatest gene probe coverage (103,511 nt captured), the highest number of unique prioritized variants, and was among the top genes for number of prioritized variants/kb (0.59).

**Table 4.** Comparing Counts of Prioritized Variants

| Gene | Unique prioritized variants | Unique patients | Gene probe coverage (nt) | Prioritized variants/patient | Prioritized variants/kb |
| --- | --- | --- | --- | --- | --- |
| *ATM* | 61 | 102 | 103511 | 0.60 | 0.59 |
| *ATP8B1* | 21 | 37 | 94793 | 0.57 | 0.22 |
| *BARD1* | 17 | 46 | 73735 | 0.37 | 0.23 |
| *BRCA1* | 19 | 24 | 52075 | 0.79 | 0.36 |
| *BRCA2* | 24 | 28 | 73332 | 0.86 | 0.33 |
| *CDH1* | 21 | 32 | 61312 | 0.66 | 0.34 |
| *CHEK2* | 12 | 13 | 28372 | 0.92 | 0.42 |
| *MLH1* | 18 | 25 | 50553 | 0.72 | 0.36 |
| *MRE11A* | 17 | 31 | 64713 | 0.55 | 0.26 |
| *MSH2* | 18 | 17 | 112437 | 1.06 | 0.16 |
| *MSH6* | 19 | 23 | 25216 | 0.83 | 0.75 |
| *MUTYH* | 8 | 16 | 21439 | 0.50 | 0.37 |
| *NBN* | 11 | 21 | 57067 | 0.52 | 0.19 |
| *PALB2* | 26 | 46 | 25319 | 0.57 | 1.03 |
| *PMS2\** | 8 | 15 | 11726 | 0.53 | 0.68 |
| *PTEN\*\** | 15 | 23 | 86059 | 0.65 | 0.17 |
| *RAD51B\*\*\** | 22 | 47 | 62465 | 0.47 | 0.35 |
| *STK11* | 12 | 20 | 28373 | 0.60 | 0.42 |
| *TP53* | 11 | 30 | 23544 | 0.37 | 0.47 |
| *XRCC2* | 7 | 10 | 19942 | 0.70 | 0.35 |

\* homologous to other genomic regions, thus fewer probes designed within gene; \*\* *PTEN* has pseudogene *PTENP1*, thus fewer probes covering exonic regions; \*\*\* probes limited to 1,000 nt surrounding all exons, and 10,000 nt up- and down-stream of gene

In total, our framework allowed for the prioritization of 346 unique variants in 246 patients, such that 85.7% of tested patients (N=287) had at least 1 prioritized variant. Most patients (84.7%) harbored fewer than 4 prioritized variants. The distribution of patients with prioritized variants was similar across eligibility groups (Table 5). Although Class 5 (91.1% of patients with prioritized variants) and Class 8 (100% with prioritized variants, with a single patient in this category) deviated to a greater extent from the mean variants/category, these differences were not significant, *χ*^2^ (4, *N*=246) = 0.98, *p* > 0.90. The distribution of prioritized variants among mutation types is: 9 protein truncating, 28 mRNA splicing, 34 affecting RBBS and/or UTR structure, 36 affecting TFBS, 119 missense and 149 affecting SRFBS, of which 29 were prioritized into multiple categories (**Supplementary Tables S14** and **S15** show this information by gene and patient, respectively).

**Table 5.** Distribution of Recruited Patients among Eligibility Groups

| Eligibility Group<sup>†</sup> | Number of Patients within Eligibility Group | Number of Patients with Prioritized Variants |
| --- | --- | --- |
| Breast cancer <60 year, and a first or second-degree relative with ovarian cancer or male breast cancer (5). | 68 | 62 |
| Breast and ovarian cancer in the same individual, or bilateral breast cancer with the first case <50 years (6). | 37 | 32 |
| Two cases of ovarian cancer, both <50 years, in first or second-degree relatives (7). | 72 | 59 |
| Two cases of ovarian cancer, any age, in first or second-degree relatives (8). | 1 | 1 |
| Three or more cases of breast or ovarian cancer at any age (10). | 109 | 92 |
|  | 287 | 246 |

The Risk Categories for Individuals Eligible for Screening for a Genetic Susceptibility to Breast or Ovarian Cancers are determined by the Ontario Ministry of Health and Long Term-Care Referral Criteria for Genetic Counseling. † Numbers in parentheses correspond to Eligibility Group designation.

All prioritized protein-truncating (N=10), and selected splicing (N=7) and missense (N=5) variants were verified by bidirectional Sanger sequencing as they were more likely to be pathogenic (taking into account available published studies). Of the protein-truncating variants, 4 nonsense, 1 indel with a conserved reading frame, and 2 frameshifts were confirmed (Table 3). Six splicing variants and all missense were confirmed (Table 1 and **Supplementary Table S13**). An additional 145 prioritized variants, including 88 non-coding variants, were confirmed upon re-sequencing of patient gDNA. Of the 57 re-sequenced coding variants, 13 were prioritized for their non-coding effects (12 SRFBS, 2 cryptic site strengthening; 1 variant prioritized for both). These variants can be found in **Supplementary Table S15** (where ‘Coverage’ column contains two or more coverage values).

### Negative Control

*ATP8B1* was sequenced and analyzed in all patients as a negative control (**Supplementary Table S16**). We prioritized 21 *ATP8B1* variants with an average of 0.22 variants/kb and 0.57 variants/patient. This is lower than the prioritization rate for many of the documented HBOC genes. This result illustrates that the proposed method represents a screening rather than a diagnostic approach, as some variants may be incorrectly prioritized.

### Pedigree Analysis

Pathogenic *BRCA2* variants within a region of exon 11 have been associated with a high incidence of OC. We therefore verified whether there were a high number of OC cases in the families of patients prioritized with exon 11 *BRCA2* variants (N=3). The family of the patient with *BRCA2* variant c.4828G>A (p.Val1610Met; diagnosed with BC at 65) has 3 reported cases of BC/OC, 1 of which is OC (diagnosed at 74), 2 degrees of separation from the proband. The patient with *BRCA2*:c.6317T>C (p.Leu2106Pro; diagnosed with BC at 52) has 3 other affected family members, 2 with OC and 1 with BC. Finally, 4 patients found to have the *BRCA2* variant c.5199C>T (p.Ser1733=) do not have any family members with reported cases of OC.

We also selected patients with prioritized mismatch repair (MMR) variants (N=8, in 10 patients) to assess the incidence of reported CRC cases in these families. Notably, the patient with mutation *MSH2*:c.1748A>G (p.Asn583Ser) had 5 relatives with CRC. A similar analysis of prioritized *CDH1* variants did not reveal any patients with a family history of gastric cancer.

### Likelihood Ratio Analyses

We carried out co-segregation analysis of 25 patients with prioritized pathogenic variants (4 nonsense, 4 frameshift, 2 in-frame deletions, 6 missense, 4 natural splicing, and 6 cryptic splicing; including a patient who exhibited prioritized natural and cryptic SS variants). We compared these findings with those from patients (N=25) harboring moderate-priority variants (variants prioritized through IT analysis only) and those in whom no variants were flagged or prioritized (N=14). In instances where disease alleles could be transmitted through either founder parent, the lineage with the highest LR was reported. For patients with likely pathogenic variants, the LRs ranged from 0.00 to 70.96 (Table 6 and **Supplementary Table S17**). Disease co-segregation was supported (LR > 1.0) in 18 patients, and the remainder were either neutral (LR < 1.0 [Mohammadi et al. 2009]) or could not be analyzed either due to missing pedigree information or limited numbers of affected individuals in a family. Patient 10-6F (*PALB2:* c.757_758delCT;) exhibited the highest likelihood (LR=70.96). Prioritized variants with neutral evidence include a variant that abolishes a natural SS in *MRE11A*, c.2070+2T>A (LR=0.03), and an in-frame deletion c.483_485delAGA in *CHEK2* (LR=0.00).

**Table 6.**
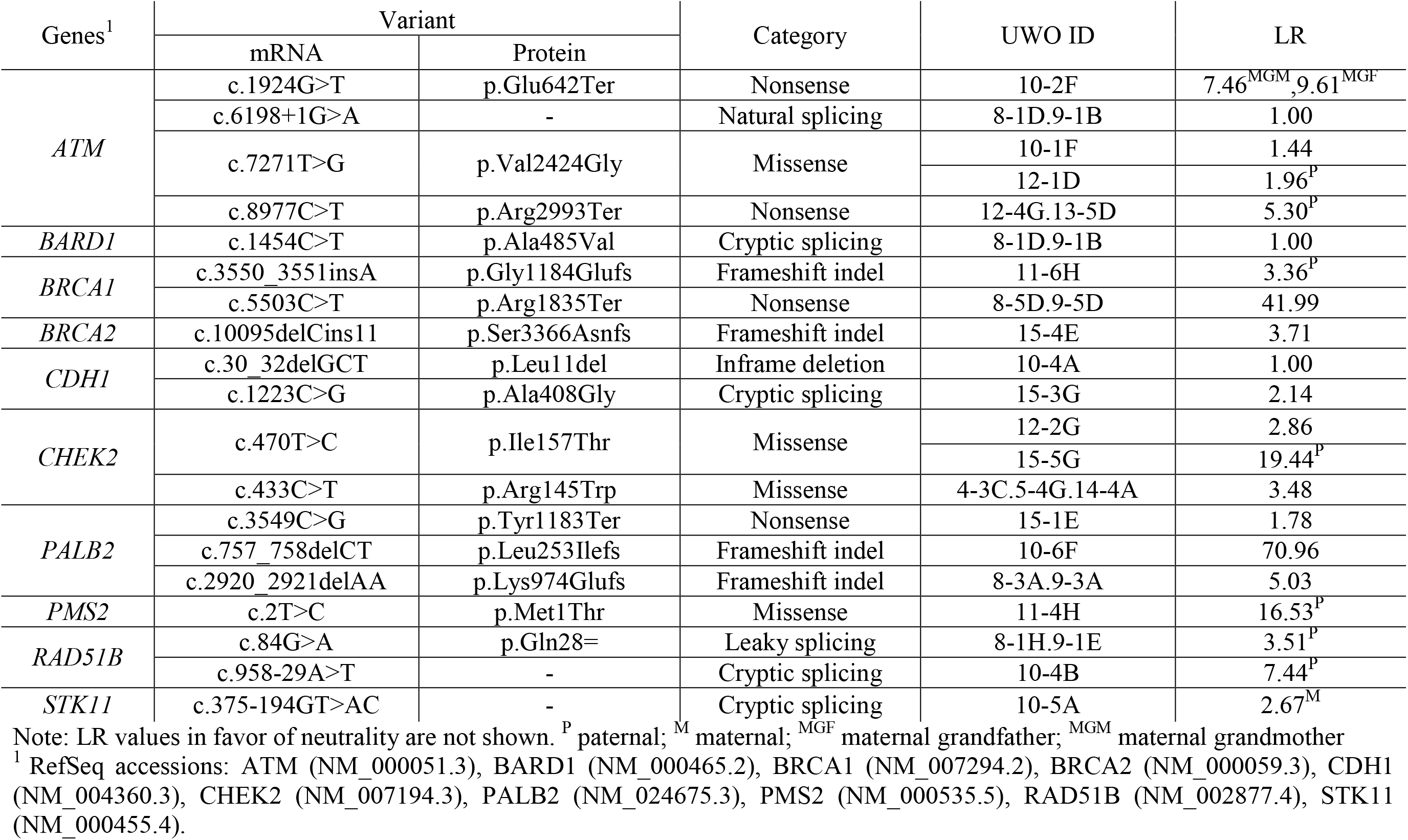
LR Values for Patients with Prioritized Truncating, Splicing, and Selected Missense Variants

## DISCUSSION

Rare non-coding and/or non-truncating mutations can confer an increased risk of disease in BC (Tavtigian et al. 2009). This study determined both coding and non-coding sequences of 20 HBOC-related genes, with the goal of discovering and prioritizing rare variants with potential effects on gene expression. This work emphasizes results from the analysis of non-coding variants, which are abundant in these genes, yet have been underrepresented in previous HBOC mutation analyses. Nevertheless, alterations to mRNA binding sites in BRCA, and lower risk or rare HBOC genes, have been shown to contribute to HBOC (ESEs in *ATM* [Heikkinen et al. 2005], *BARD1* [Ratajska et al. 2011], and *BRCA* genes [Gochhait et al. 2007; Sanz et al. 2010]). We prioritized 346 unique variants that were predicted to result in 4 nonsense, 3 frameshift, 2 indels with preserved reading frame, 119 missense, 4 natural splicing, 6 cryptic splicing, 17 pseudoexon activating, 148 SRFBS, 36 TFBS, 3 UTR structure, and 31 RBBS mutations (**Supplementary Table S14**). Among these variants, 101 were novel (see **Supplementary Table S18** for references to previously identified variants). Compared to our initial 7-gene panel (Mucaki et al. submitted), the inclusion of the additional genes in this study prioritized at least 1 variant in 15% additional patients (increased from 70.6 to 85.7%).

The *BRCA* genes harbor the majority of known germline pathogenic variants for HBOC families (Chong et al. 2014). However, a large proportion of the potentially pathogenic variants identified in our study were detected in *ATM*, *PALB2*, and *CHEK2*, which although of lower penetrance, were enriched because the eligibility criteria excluded known *BRCA1* and *BRCA2* carriers. *BRCA1* and *BRCA2* variants were nevertheless prioritized in some individuals. We also had expected intragenic clustering of some *BRCA* coding variants (Mucaki et al. 2011). For example, pathogenic variants occurring within exon 11 of *BRCA2* are known to be associated with higher rates of OC in their families (Lubinski et al. 2004). We identified 3 variants in exon 11; however there was no evidence of OC in these families. Overall, *ATM* and *PALB2* had the highest number of prioritized variants (61 and 26, respectively). However, only 12 variants were prioritized in *CHEK2;* potentially pathogenic variants may have been under-represented during sequence alignment as a consequence of the known paralogy with *CHEK2P2*.

Fewer *TP53, STK11*, and *PTEN* variants were prioritized, as pathogenic variants in these genes tend to be infrequent in patients who do not fulfill the clinical criteria for their associated syndromes (Li-Fraumeni syndrome, Peutz-Jeghers syndrome, and Cowden syndrome, respectively [Hollestelle et al. 2010]) although they have been indicated as near moderate to high-risk genes in breast cancer (Easton et al. 2015). This underrepresentation of prioritized variants may be supported by the negative Residual Variation Intolerance Scores (RVIS) for these genes (Petrovski et al. 2013), which are likely indicative of purifying selection. Although the density of prioritized variants in these genes is below average (18.2 per gene), the total number was nonetheless important (*TP53*=11, *STK11*=12, *PTEN*=15).

The fundamental difference between IT and other approaches such as CADD (Combined Annotation Dependent Depletion; Kircher et al. 2014) is that IT depends only on positive experimental data from the same or closely related species. CADD doesn’t appear to account for unobserved reversions or other hidden mutations (eg. perform a Jukes-Cantor correction; Jukes and Cantor, 1969), nor are the effects of these simulated. Furthermore, the CADD scoring system is *ad hoc*, which contrasts with strong theoretical basis on the information theory approach (Rogan and Schneider, 1995) in which information changes in bits represent a formally proven relationship to thermodynamic stability, and therefore can be used to measure binding affinity. This makes it different from other unitless methods with unknown distributions, in which differences binding affinity cannot be accurately extrapolated from derived scores.

We compared the frequency of all prioritized variants in our patient cohort to the population allele frequencies (1000 Genomes Project, Phase 3; http://www.1000genomes.org; 1000 Genomes Project Consortium, 2012) to determine if variants more common in our cohort might be suggestive of HBOC association. Three variants in at least 5 HBOC patients are present at a much lower frequency in the general population than in our HBOC population. *NBN* c.*2129G>T, present in 4.18% of study cohort, is considerably rarer globally (0.38% in 1000Genomes; < 0.1% in other populations). Similarly, the *RAD51B* c.-3077G>T variant (2.09%), is rare in the general population (0.08%). Interestingly, *BARD1* c.33G>T (1.74% of study cohort) has only been reported in the American and European populations in 1000Genomes (0.29% and 0.20%, respectively) and only Europeans in the Exome Variant Server (0.24%; http://evs.gs.washington.edu/EVS/). In Southwestern Ontario, individuals are often of American or European ancestry. The variant was found to be more common in the Exome Aggregation Consortium (ExAC; http://exac.broadinstitute.org/) in 1.17% tested Finnish population (0.41% in their non-Finnish European cohort), though no alleles were found in the Finnish populations in 1000 Genomes (N=99). Therefore, the allele frequency of this *BARD1* variant in our HBOC population may simply be enriched in a founder subset of general populations. While we cannot rule out skewing of these allele frequencies due to population stratification, our findings suggest that gene expression levels could be impacted by these variants.

We applied sub-populations allele frequency analysis for all of our prioritized variants. **Supplementary Table S19** lists the 49 variants that have allele frequencies > 1% in various sub-population (based on dbSNP). Allele frequencies were as high as 4.2% for the *BRCA2* c.40+192C>T (8-1G.9-1C), predicted to affect transcription factor binding, in the East Asian sub-population. Without additional information on patient ethnicities, it is not possible to eliminate prioritized variants that are common in specific sub-populations.

Co-segregation analysis is recommended by the American College of Medical Genetics and Genomics (ACMG) for variant classification (Richards et al. 2015). Among patients with likely pathogenic, highly penetrant mutations in our cohort (N=24), some variants had LR values consistent with causality, whereas others provided little evidence to support co-segregation among family members (Table 6 and **Supplementary Table S17**). An important caveat however was that use of *BRCA2* penetrance values in non*-BRCA* genes may have resulted in underestimates of LR values.

In order to evaluate the application of co-segregation analysis in the context of this study, we chose to perform this analysis on patients with moderate priority variants (i.e. variants affecting binding sites) and patients with no flagged or prioritized variants (N=25 and 14, respectively). LRs ranged from 0.0034 to 78.0 for moderate-priority variants and from 0.0005 to 57.0 for patients with no flagged or prioritized variants (**Supplementary Figure S2**). The proportion of LR values supporting neutrality and those supporting causation was comparable between patients with prioritized, moderately prioritized, and flagged variants (**Supplementary Figure S2**). This suggests co-segregation analysis is only useful in the context of other supporting results for assessing pathogenicity (eg. likelihood of being pathogenic or benign). Furthermore, the lack of genotype information and at times smaller pedigrees likely also contributed to the lack of concordance between LRs and variant priority.

A small number of patients with a known pathogenic variant carried other prioritized variants. These were likely benign or possibly, phenotypic modifiers. One patient possessed 5 prioritized variants (1 missense, 1 SRFBS, 1 TFBS, and 2 RBBSs) in addition to a *BRCA1* nonsense mutation (c.5503C>T). While these variants may not directly contribute to causing HBOC, they may act as a risk modifier and alter expression levels (Antoniou and Easton 2006).

Similarly, genes lacking association with HBOC can be used as a metric for determining a false-positive rate of variant prioritization. In this study, we prioritized 21 *ATP8B1* variants among 37 of our HBOC patients (**Supplementary Table S16**) despite it having not been previously associated with any type of cancer. A variant with a deleterious effect on *ATP8B1* may lead to *ATP8B1*-related diseases, such as progressive familial intrahepatic cholestasis (Gonzales et al. 2014), but should not increase the chances of developing BC. Thus, while our framework may be effective at prioritizing variants, only genes with previous association to a disease should be included in analyses similar to the present study to minimize falsely prioritized variants.

Additional workup of prioritized non-coding and non*-BRCA* variants is particularly important, because with few exceptions (Easton et al. 2015), the pathogenicity of many of the genes and variants has not been firmly established. Furthermore, mutations in several of these genes confer risk to other types of cancer, which alters the management of these patients (Knappskog and Lønning 2012). The next step towards understanding the role these prioritized variants play in HBOC is to test family members of the proband and to carry out functional analysis. If this is not possible, then their effects on gene expression could be evaluated using assays for RNA stability and RNA localization. Protein function could be evaluated by binding site assays, protein activity, and quantitative PCR.

A significant challenge associated with VUS analysis, particularly in the case of many of these recent HBOC gene candidates, is the under-reporting of variants and thus positive findings tend to be over-represented in the literature (Kraft 2008). Hollestelle et al. (2010) argue that a more stringent statistical standard must be applied (i.e. *p*-values of 0.01 should be used as opposed to 0.05) to under-reported variants (namely in moderate-risk alleles), because of failure to replicate pathogenic variants, which we have also found (Viner et al. 2014). In the same way that we use IT-based analysis to justify prioritizing variants for further investigation, variants that are disregarded as lower priority (and that are likely not disease-causing) have been subjected to the same thresholds and criteria. Integrating this set of labeled prioritized and flagged, often rare variants, from this cohort of *BRCA*-negative HBOC patients, to findings from exome or gene panel studies of HBOC families should accelerate the classification of some VUS.

Different variant interpretation and reporting guidelines consider the reporting of VUS to be either optional or essential (Wallis et al. 2013; Richards et al. 2015). In all cases, a reported VUS cannot be the basis for a clinical decision and should be followed up and further investigated. In any case, the number of reported VUS in an individual are frequently too extensive for detailed characterization. Reducing the full set of variants obtained by complete gene sequencing to a prioritized list will be an essential prerequisite for targeting potentially clinically relevant information. Informing patients of prioritized VUS may increase patient accrual and participation (Murphy et al. 2008). However, it will be critical to explain both the implications and significance of prioritization and the limitations, namely counselling patients to avoid clinical decisions, based on this information (Vos et al. 2012).

## Acknowledgements

We would like to thank the patients who participated, Karen Panabaker for oversight in assessing patient eligibility, directing patient recruitment, and follow-up counseling, and Dr. Peter Ainsworth and Alan Stuart for access to genomic DNA samples and robotic equipment in the MGL at London Health Sciences Centre. Abby Watts-Dickens assisted in patient recruitment. PKR is supported by the Canadian Breast Cancer Foundation, Canadian Foundation for Innovation, Canada Research Chairs Secretariat, and the Natural Sciences and Engineering Research Council of Canada (NSERC Discovery Grant 371758-2009). NGC was supported by CIHR Strategic Training Program in Cancer Research and Technology Transfer and the Pamela Greenaway-Kohlmeier Translational Breast Cancer Research Unit. Access to the Shared Hierarchical Academic Research Computing Network (SHARCNET) and Compute/Calcul Canada is gratefully acknowledged.

